# Systemic hypoxia drives glycogen-fueled progression of lung adenocarcinoma

**DOI:** 10.64898/2026.09.01.748693

**Authors:** Harrison A. Clarke, Cameron J. Shedlock, Haley Peters, Tara R. Hawkinson, Damon D. Kooser, Olivia Janzen, Ariya Varma, Franca Bucco, Charles M. Soto, Reece Larson, Alison M. Ryan, Terrymar Medina, Roberto A. Ribas, Christopher M. Calulot, Lei Wu, Jenny Russ, Sophia Florea, Yanfei Wang, Kari B. Basso, Derek B. Allison, Manuela Corti, David D. Fuller, Warren J. Alilain, Gordon S. Mitchell, Barry J. Byrne, Craig W. Vander Kooi, Yi Guo, Matthew S. Gentry, Ramon C. Sun

## Abstract

In advanced stages, lung adenocarcinoma obstructs airways and disrupts ventilation-perfusion relationships in the lung, causing systemic hypoxemia and enabling a feed-forward loop that accelerates malignancy. Systemic hypoxemia is also experienced due to common respiratory comorbidities such as chronic obstructive pulmonary disease (COPD) and obstructive sleep apnea (OSA), potentially accelerating malignancy. In a statewide electronic health record network, pre-existing COPD (598 matched pairs) or sleep apnea (235 matched pairs) independently predicted worse survival following incident lung cancer diagnosis. Since the mechanistic basis of the link between malignancy and hypoxia is not well understood, we created systemic hypoxia in Kras ^LSL-G12D/+^;Trp53^fl/fl^ (KP) mice by delivering low inspired oxygen concentrations (8% inspired oxygen; 8 h daily). Hypoxia nearly doubled tumor multiplicity and selectively remodeled cancer central carbon metabolism. Spatially resolved metabolomics revealed marked tumor-compartment glycogen accumulation, elevated tricarboxylic-acid cycle intermediates, and depleted glycolytic pools. Quantitative proteomics across cellular models and autochthonous tumors demonstrated that systemic hypoxia drives glycogen mobilization selectively through the lysosomal enzyme acid α-glucosidase (GAA). Tumor-cell-autonomous deletion of GAA eliminated the hypoxia-driven growth advantage and disrupted downstream anabolic biosynthetic pathways. Thus, systemic hypoxia drives lung adenocarcinoma expansion by mobilizing lysosomal glycogen reserves through GAA to sustain proliferative growth.

**Highlights:**

- Systemic hypoxia acts as an intrinsic driver of lung adenocarcinoma progression.
- Controlled systemic hypoxia doubles tumor multiplicity and reprograms cancer metabolism.
- Hypoxic tumors mobilize glycogen via lysosomal GAA independently of HIF-1α stabilization.
- Tumor-cell-autonomous deletion of GAA eliminates the hypoxia-mediated growth advantage in vivo.
- Pre-existing COPD and sleep apnea exacerbate hypoxemic burden and worsen clinical survival.

**eTOC Blurb:** Lung tumors compromise pulmonary gas exchange, driving systemic hypoxemia that accelerates malignancy. Comorbidities such as chronic obstructive pulmonary disease (COPD) and obstructive sleep apnea compound this hypoxemic burden. Clarke et al. isolate systemic hypoxia in mice, showing it doubles lung tumor multiplicity by inducing glycogen degradation through lysosomal acid α-glucosidase (GAA). Deleting GAA eliminates this hypoxic growth advantage.

## Introduction

Lung adenocarcinoma is a leading cause of cancer death worldwide^1^. As lung tumors expand, they infiltrate healthy lung parenchyma, obstruct conducting airways, and collapse functional alveolar space (*i.e.,* atelectasis). This progressive structural damage leads to severe ventilation-perfusion mismatch, impaired gas exchange and hypoxemia^2^. Consequently, hypoxemia is an intrinsic product of the expanding tumor mass. While reduced oxygen delivery stresses tissues host-wide, localized low oxygen within the tumor microenvironment accelerates cellular proliferation, invasiveness, and treatment resistance^3^. Since tumor mass and systemic hypoxemia progress in tandem, it is difficult to decouple the biological impact of systemic hypoxemia from local airway effects without an experimental animal model.

Common respiratory comorbidities significantly compound this baseline hypoxic burden. Chronic obstructive pulmonary disease (COPD) and obstructive sleep apnea (OSA) represent the most prevalent chronic respiratory comorbidities in lung cancer patients^4,5^. COPD increases lung cancer incidence up to six-fold (independent of smoking history)^6^, and sleep apnea increases cancer incidence in millions of individuals^7^. Although COPD impairs gas exchange through maldistribution of ventilation and perfusion within the lung^8^, sleep apnea is caused by repetitive upper-airway collapse and episodic hypoxia during the ensuing apneas^9^. Despite these distinct mechanisms, both disorders converge on the same consequence: a fall in systemic oxygenation ^10^. Thus, whereas advanced COPD imposes sustained hypoxemia across the day and night^10^, severe sleep apnea causes hundreds of transient desaturations each night, increasing the cumulative nocturnal hypoxic burden ^10^. Although cancer mortality scales directly with this nocturnal hypoxic burden^11,12^, it does not scale with the frequency of airway occlusions^13^. Systemic hypoxia is the shared insult linking these respiratory disorders to shortened patient survival. However, clinical datasets cannot easily isolate hypoxia from confounding variables such as cigarette-smoke toxins and chronic airway inflammation. Because isolating systemic hypoxia in the context of lung tumorigenesis has not been previously done, testing it as a pure independent variable requires a highly controlled experimental model. To achieve this, we modeled systemic hypoxemia in rodents by delivering a defined gas mixture with significantly lowered oxygen inside a chamber.

The next step was to understand how this hypoxemia alters tumor metabolism using advanced spatial profiling. Lung adenocarcinoma maintains a robust oxidative metabolism. However, instead of executing a simple glycolytic switch, human tumors oxidize glucose through the tricarboxylic-acid cycle^14^. Likewise, in Kras-driven mouse lung tumors, growth depends on glucose oxidation through the tricarboxylic-acid cycle, and these tumors form and grow normally even without hypoxia-inducible factor 1-alpha (HIF-1α)^15,16^. To sustain this oxidative flux under metabolic stress, these tumors require a flexible metabolic reservoir. Glycogen represents an established oncogenic driver in lung adenocarcinoma^17^, accumulating across diverse driver mutations to track tumor histological grade, predict poor patient survival, and accelerate tumorigenesis^17^. Aberrant glycogen metabolism also drives other malignancies, including hepatocellular carcinoma^18^, breast cancer^19^, ovarian clear-cell carcinoma^20^, and Ewing sarcoma^21^. Cancer cells typically degrade glycogen through two distinct biochemical routes. Cytosolic glycogen phosphorylase releases glucose-1-phosphate to support glycolysis in near-anoxic microenvironments^22,23^, whereas lysosomal acid α-glucosidase (GAA) hydrolyzes glycogen to free glucose downstream of nutrient and oxygen stress^24,25^. Whether systemic hypoxemia mobilizes lung tumor glycogen remains unknown, and the specific catabolic enzyme engaged under this systemic stress is unresolved. Because glycogen utilization is highly context-specific across different tumor types^19,26^, defining the exact enzymatic route in lung cancer is necessary to identify a targetable metabolic vulnerability.

To address this, we used a controlled environmental chamber to impose systemic hypoxemia in vivo. We exposed mice to a daily 8-hour block of 8% inspired oxygen followed by 16 hours of room air, a regimen that models daily systemic hypoxemia while removing other variables such as smoking, infection, and treatment confounders. We tested hypoxia effects in 4 complementary systems: 1) we analyzed statewide electronic health records, 2) mapped metabolism in wild-type and KP mouse models, 3) applied spatial MALDI mass spectrometry imaging to resected human tumors, and 4) built *in vitro* kinetic flux models. Finally, we tested the functional requirements of this metabolic pathway *via* genetic loss of GAA in autochthonous lung tumors. Ultimately, we demonstrate that systemic hypoxia accelerates lung adenocarcinoma progression by driving GAA-dependent lysosomal glycogen catabolism independently of the canonical HIF-1α program.

## Results

### Pre-existing respiratory comorbidity is associated with worse survival in lung cancer

Because direct blood oxygen measurements are rarely captured in real-world electronic health records, we investigated chronic obstructive pulmonary disease (COPD) and obstructive sleep apnea as clinical proxies for hypoxemia, under a working model in which impaired pulmonary oxygen uptake lowers arterial oxygen content (Figure 1A). To evaluate whether these pre-existing comorbidities correlate with reduced lung cancer survival, we constructed two independent propensity score-matched cohorts using the statewide OneFlorida+ network (Methods). Patients carrying a pre-existing COPD diagnosis exhibited significantly worse survival compared to their matched controls, recording 199 deaths among 598 patients with COPD versus 165 deaths among 598 controls. This corresponded to an HR for all-cause mortality of 1.44 (95% CI: 1.17–1.78), with survival curves separating within the first year and remaining separated through five years (Figure 1B). Kaplan-Meier estimates revealed cumulative mortalities of 27.2% versus 16.9% at one year, 42.5% versus 33.7% at three years, and 48.7% versus 40.7% at five years (Figure 1C). This survival disadvantage persisted across both sexes and remained stable across alternative covariate-adjusted models.

**Figure 1.**
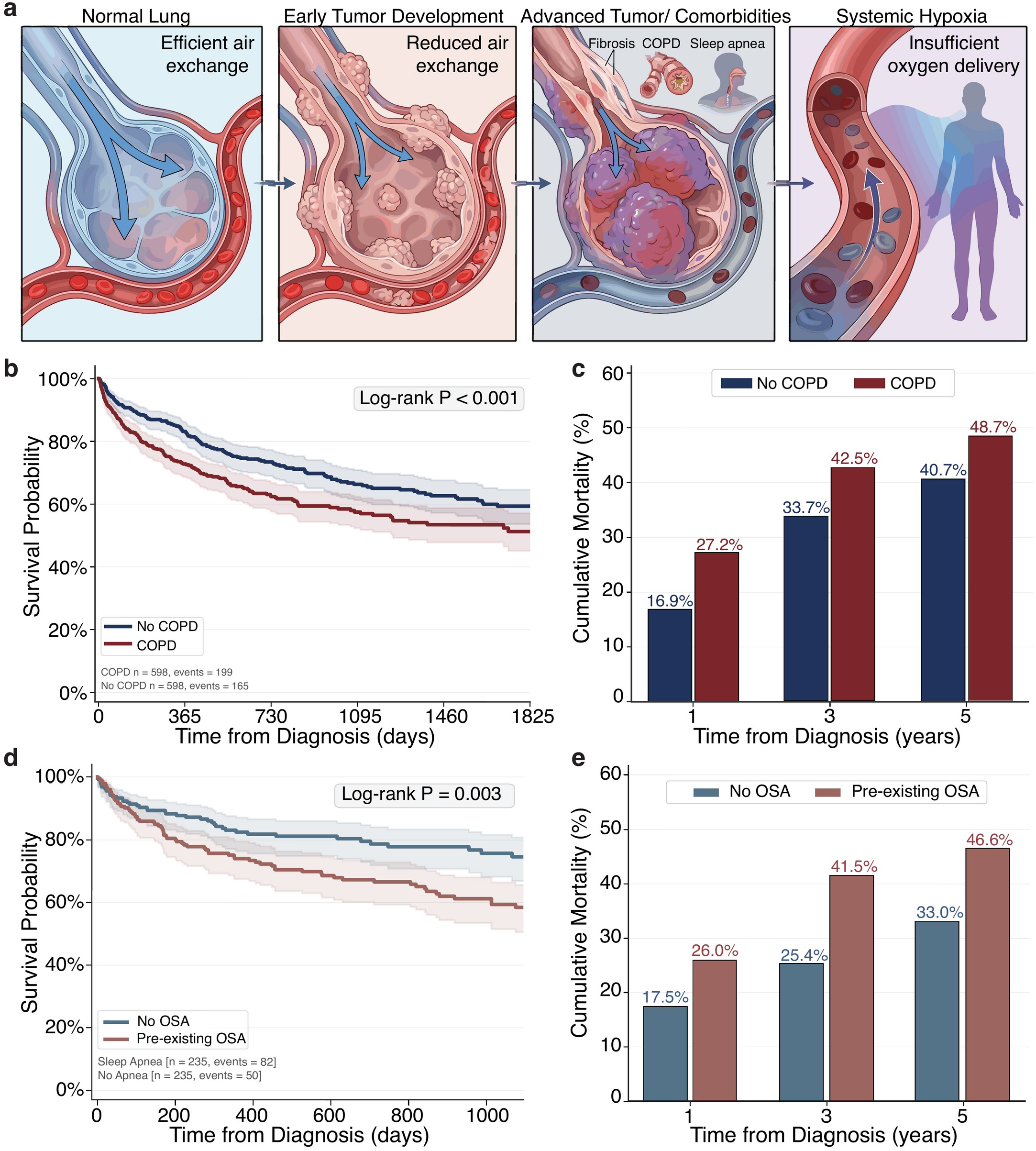
Pre-existing COPD or obstructive sleep apnea is associated with worse survival after lung cancer diagnosis. (a) Schematic illustrating the relationship between advancing pulmonary pathology, respiratory comorbidities (pulmonary fibrosis, COPD, or obstructive sleep apnea [OSA]), and impaired systemic oxygen delivery. (b and c) Kaplan-Meier survival curves (b) and cumulative mortality rates at 1, 3, and 5 years (c) for the matched COPD cohort (n = 598 patients per group; 199 deaths among COPD patients versus 165 deaths among matched controls; log-rank p < 0.001). Absolute mortality differences were 10.3% (1 year), 8.8% (3 years), and 8.0% (5 years). (d and e) Kaplan-Meier survival curves (d) and cumulative mortality rates at 1, 3, and 5 years (e) for the matched OSA cohort (n = 235 patients per group; 82 deaths among OSA patients versus 50 deaths among matched controls; log-rank p = 0.003). Absolute mortality differences were 8.5% (1 year), 16.1% (3 years), and 13.6% (5 years). Statistics: shaded bands represent 95% confidence intervals (CIs). Cohorts were propensity-score matched 1:1 with exact matching on cancer stage. Hazard ratios (HR) were calculated using Cox proportional-hazards regression: COPD HR 1.44 (95% CI: 1.17–1.78; p = 0.0006); OSA HR 1.70 (95% CI: 1.19–2.42; p = 0.003).

We observed a similarly detrimental effect in the sleep apnea cohort, where 82 deaths occurred among 235 patients compared to 50 among 235 matched controls. This yielded an HR of 1.70 (95% CI: 1.19–2.42; Figure 1D), with cumulative mortality reaching 26.0% versus 17.5% at one year, 41.5% versus 25.4% at three years, and 46.6% versus 33.0% at five years (Figure 1E). While each estimate was derived within a propensity score-matched cohort, both cohorts rely on diagnostic billing codes without objective oximetry or disease severity metrics, precluding us from confirming active hypoxemia in specific patients. Furthermore, retrospective registries inherently harbor residual confounding, such as varying smoking intensities and unmeasured cancer-directed therapies. These clinical signatures suggest that systemic hypoxia contributes to poor survival, but they cannot definitively prove causation. To test our hypothesis that systemic hypoxia alone is biologically sufficient to accelerate lung tumor growth, we transitioned to a controlled in vivo animal model.

### Systemic hypoxia preserves normal lung function and the baseline metabolome while selectively expanding glycogen stores

To isolate the biological effect of systemic hypoxemia in vivo, we exposed wild-type mice to a daily eight-hour block of 8% inspired oxygen over two weeks (Figure 2A). We first measured ventilatory parameters using whole-body plethysmography to confirm that the animals could physiologically tolerate this chronic regimen. Minute ventilation was preserved across the two-week exposure, remaining indistinguishable from the pre-exposure baseline at both day 7 and day 14 (day 14, 4.02 versus 4.18 mL/min/g; n = 8; Figures 2B and 2C; see Figure S2). The mice achieved this homeostatic adaptation by breathing more deeply and slowly; while frequency remained lowered (269 versus 366 breaths/min; 0.74-fold), tidal volume significantly increased from 0.0107 to 0.0150 mL/g (1.40-fold)^27,28^. Hypercapnic-hypoxic chemoreflex sensitivity also remained fully preserved on day 14 (1.75-fold versus 1.81-fold baseline increase; see Figure S1).

**Figure 2.**
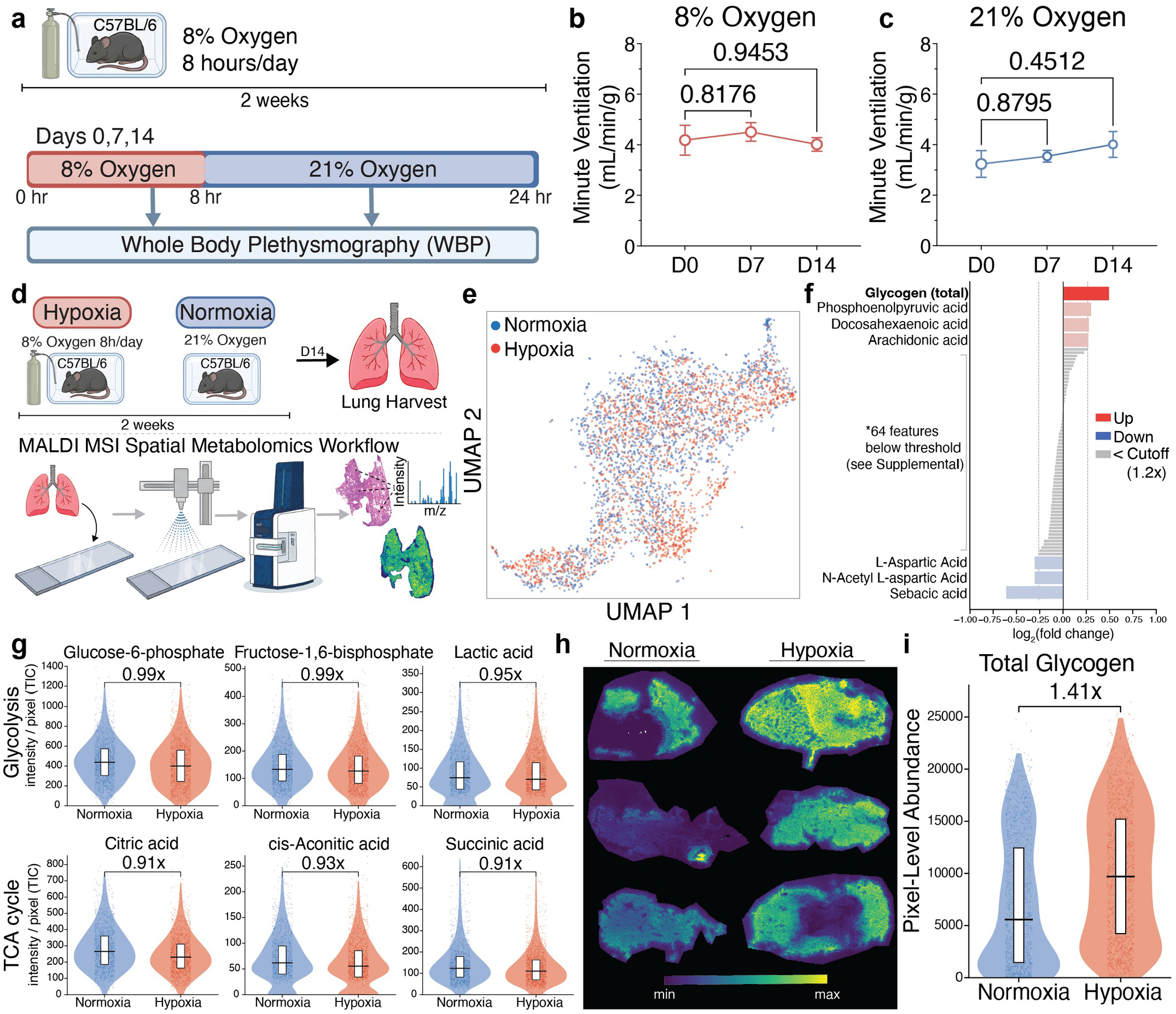
Chronic systemic hypoxia selectively raises lung glycogen without impairing ventilation. (a) Experimental schedule for intermittent hypoxia exposure (8% oxygen, 8 h/day during rest phase for 2 weeks) and whole-body plethysmography in C57BL/6 mice. (b and c) Minute ventilation normalized to body weight measured during the 8% oxygen block (b) and 21% oxygen block (c) at baseline (D0), day 7 (D7), and day 14 (D14) (n = 8 mice per group). Data are mean ± SEM. (d) Workflow for spatial MALDI mass spectrometry imaging on tumor-free lungs harvested at day 14 from normoxic and hypoxic wild-type mice (n = 5 mice per group). (e) UMAP projection of individual tissue pixels colored by exposure condition. (f) Ranked log2 fold change (hypoxia versus normoxia) for all 71 detected metabolic features (n = 5 mice per group). Red indicates increased, blue indicates decreased (≥1.2-fold cutoff), and gray indicates unchanged features. (g) Pixel-level intensity distributions for representative glycolytic and tricarboxylic-acid (TCA) cycle intermediates (n = 5 mice per group). Violins show pixel distributions, white boxes indicate interquartile ranges, and horizontal bars denote medians. (h and i) Representative glycogen ion images (h) and total pixel-level abundance summed across 17 chain-length channels (i) (n = 5 mice per group; 136,518 normoxic and 180,645 hypoxic pixels; 1.41-fold per-mouse mean increase). Statistics: metabolite intensities were total ion count (TIC)-normalized and log10-transformed. Pixel comparisons were performed by two-tailed Student’s t test with Benjamini-Hochberg false discovery rate (FDR) correction; biological significance is defined by a ≥1.2-fold change. See also Figures S1–S3.

Because the mice functionally adapted to this exposure, we asked whether their baseline non-malignant lung metabolome remained similarly stable. Using spatial MALDI mass spectrometry imaging on tumor-free lungs (n = 5; Figure 2D), we found that 64 of 71 detected metabolites were entirely unchanged at the end of the two-week challenge (Figure 2F; see Figure S3). Unsupervised principal component analysis confirmed this stability, showing no distinct separation between normoxic and hypoxic parenchyma (Figure 2E). Core glycolytic intermediates, tricarboxylic-acid cycle metabolites, adenine nucleotides, and the glutathione redox couple remained constant (Figure 2G), with only minor fluctuations observed in species like phosphoenolpyruvate (1.23-fold) and aspartate (0.83-fold). Against this remarkably stable metabolic background, glycogen emerged as the singular expanded carbon pool, increasing 1.41-fold across the entire chain-length ladder and localizing to discrete parenchymal deposits (Figures 2F, 2H, and 2I). Thus, systemic hypoxia selectively expands normal lung glycogen stores without disrupting broader carbon metabolism or ventilatory capacity.

### Systemic hypoxia increases tumor burden and remodels cancer metabolism

Having defined this stable physiological baseline in wild-type mice, we next tested how systemic hypoxia alters cancer progression using the autochthonous Kras^LSL-G12D/+^;Trp53^fl/fl^ (KP) model^17,29,30^. To specifically isolate tumor progression rather than initiation, we allowed adenoviral Cre-initiated tumors to establish for 30 days before applying the hypoxic exposure from day 30 to day 100 (Figure 3A). In stark contrast to the resilient normal lung tissue, the tumor compartment was profoundly vulnerable to systemic hypoxemia. Histological analysis revealed visibly larger tumor areas in the hypoxic lungs (Figure 3B), and quantitative assessment confirmed that tumor multiplicity nearly doubled, rising from 22.7 to 45.0 tumors per lung (1.99-fold increase; n = 6/5; Figure 3C). Systemic hypoxia therefore actively accelerates lung adenocarcinoma progression in vivo.

**Figure 3.**
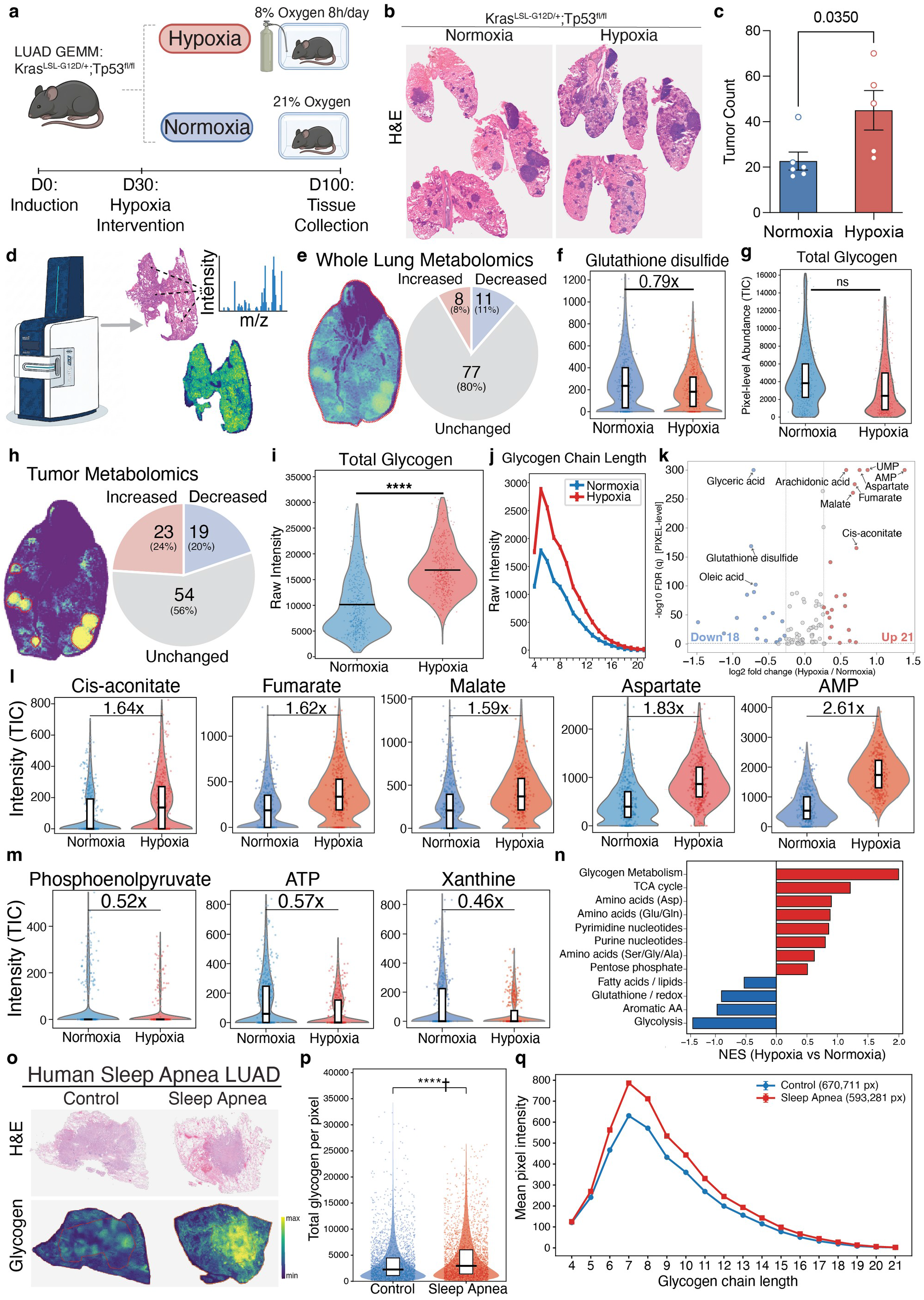
Spatial metabolomics reveals glycogen accumulation within hypoxic lung tumors and in human sleep-apnea non-small cell lung cancer. (a and b) Experimental schema for adenoviral Cre induction in Kras^LSL-G12D/+^;Trp53^fl/fl^ (KP) mice under normoxia or 8% oxygen (a) and representative hematoxylin and eosin (H&E) histology (b). (c) Histological tumor counts per lung (n = 6 normoxic, 5 hypoxic mice; two-tailed Student’s t test, p = 0.035). Data are mean ± SEM with individual biological replicates overlaid. (d) MALDI imaging segmentation workflow isolating tumor regions from adjacent non-malignant parenchyma. (e–g) Whole-lung metabolic profiling across 96 small molecules (e), glutathione disulfide (f), and total glycogen (g) (n = 9 normoxic and 6 hypoxic tissue pieces from 3 and 2 mice, respectively). (h–m) Metabolic profiling strictly within the segmented tumor compartment (n = 5 tumors per group, each from an independent animal). Shown are the tumor small-molecule response (h), total raw glycogen intensity (i), glycogen chain-length distribution (degree of polymerization [DP] 4–21) (j), volcano plot highlighting 21 increased and 18 decreased metabolites (q < 0.1, ≥1.2-fold) (k), elevated TCA intermediates and AMP (l), and depleted glycolytic/high-energy intermediates (m). (n) Metabolite-set enrichment analysis (weighted Kolmogorov-Smirnov on signed t rankings, 2,000 permutations, Benjamini-Hochberg correction). (o–q) Matched H&E and glycogen ion images (o), total glycogen quantification (p), and chain-length distribution (q) in human non-small cell lung cancer resected from patients with or without pre-existing sleep apnea (n = 5 patients per group, matched for stage and grade; 670,711 control and 593,281 sleep-apnea pixels). Statistics: MALDI data were TIC-normalized (except raw glycogen) and evaluated at pixel level with Benjamini-Hochberg correction; biological significance is defined by a ≥1.2-fold change. Human tissues were acquired without normalization. See also Figures S4 and S5, and Table S1.

To understand the metabolic mechanism driving this accelerated growth, we profiled the tumor-bearing lungs. Because dense normal parenchyma dilutes the tumor signal, whole-lung tissue homogenates masked these cancer-specific shifts. Across the whole lung, 77 of 96 quantified metabolites were unchanged (Figures 3E and 3F; see Figure S4), and total glycogen actually trended downward (0.78-fold; Figure 3G). To overcome this dilution, we applied spatial segmentation to isolate solid-tumor pixels from the adjacent normal tissue (Figure 3D)^21,31^. Within this isolated tumor compartment, the metabolic response was profound: 23 metabolites rose and 19 fell beyond the 1.2-fold cutoff (Figure 3H; see Figure S5). Tumor glycogen was significantly elevated under systemic hypoxia across the entire chain-length ladder, particularly among longer chains (Figures 3I and 3J). Systemic hypoxia comprehensively reprogrammed tumor central-carbon metabolism, with 21 metabolites significantly increased and 18 decreased (Figure 3K). Tricarboxylic-acid cycle intermediates spiked, including cis-aconitate (1.64-fold), fumarate (1.62-fold), malate (1.59-fold), and aspartate (1.83-fold; Figure 3L). Conversely, high-energy phosphates and glycolytic intermediates dropped, including phosphoenolpyruvate (0.52-fold), ATP (0.57-fold), and xanthine (0.46-fold; Figure 3M). Several metabolites shifted in opposite directions between the tumor and adjacent parenchyma^31^. Metabolite-set enrichment analysis confirmed these dramatic shifts, ranking glycogen metabolism as the top upregulated pathway (NES 2.00), followed by the tricarboxylic-acid cycle (1.21), with glycolysis emerging as the most depleted set (−1.37; Figure 3N).

We previously established that aberrant glycogen accumulation is already a baseline phenotype of both human and mouse lung adenocarcinoma^17^. To test whether a hypoxemic comorbidity further exacerbates this accumulation as observed in our mouse model, we examined resected tumors from patients with (n = 5) or without (n = 5) a pre-existing diagnosis of obstructive sleep apnea (Table S1). We carefully matched these clinical specimens for stage and histological grade to ensure comparability. Spatial MALDI imaging revealed visibly elevated glycogen within the tumors resected from patients with sleep apnea (Figure 3O), and quantitative analysis confirmed that total glycogen was 1.21-fold higher in this cohort (4,542 versus 3,755 arbitrary units; Figure 3P). While the glycogen chain-length distribution peaked at a chain length of seven in both groups, tumors from patients with sleep apnea showed consistent, broad elevations across chain lengths five through thirteen (Figure 3Q). Collectively, these data provide evidence that systemic hypoxia selectively remodels the tumor compartment to elevate oxidative flux and further expand glycogen reserves.

### Hypoxic cancer cells dynamically mobilize glycogen to fuel anabolic metabolism

To isolate the direct effect of 8% hypoxia on cancer cell metabolism over time, we established a controlled in vitro system. We cultured human A549 lung adenocarcinoma cells under 8% gas-phase oxygen or standard normoxia, and we profiled their metabolomes using MALDI mass spectrometry imaging at 8 and 24 hours (n = 8; Figure 4A).

**Figure 4.**
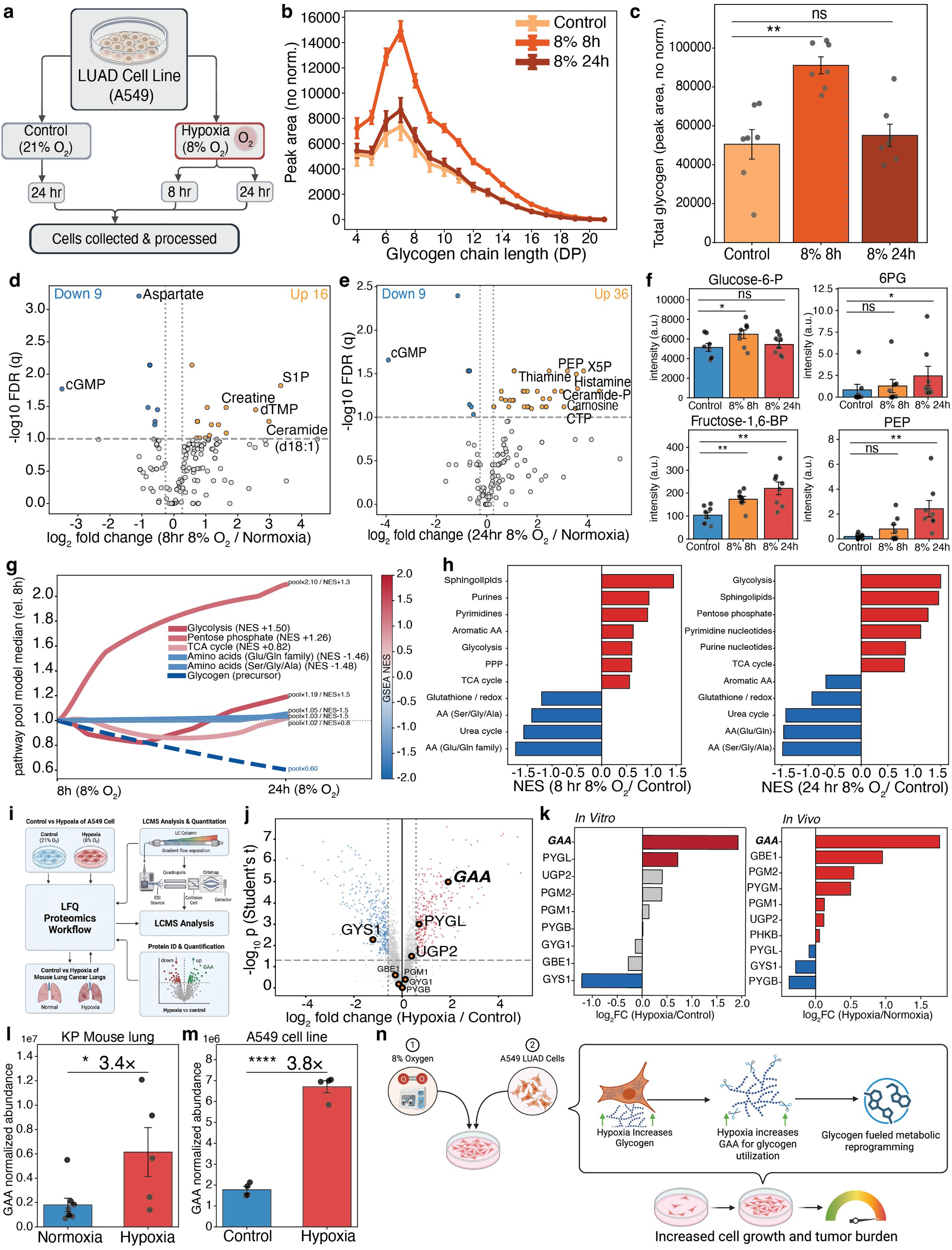
Hypoxic lung adenocarcinoma cells dynamically mobilize glycogen and selectively induce lysosomal GAA. (a) Design of the A549 hypoxia time course (21% oxygen control versus 8% oxygen for 8 or 24 h; n = 8 regions per group for metabolomics, n = 7–8 for glycogen, n = 4 for proteomics). (b and c) Glycogen chain-length distribution (DP 4–21) (b) and total glycogen abundance (c) under hypoxia. Data are mean ± SEM. (d and e) Volcano plots of the A549 metabolome at 8 h (d) and 24 h (e) of hypoxia compared to normoxia (region-level Student’s t test with Benjamini-Hochberg correction; q < 0.1, ≥1.2-fold). (f) Intensity of representative glycolytic and pentose phosphate pathway intermediates. Data are mean ± SEM with individual regions overlaid. (g) Kinetic metabolic network model integrated from the 8 h to 24 h hypoxic state. (h) Metabolite-set enrichment analysis at 8 h and 24 h relative to control (weighted Kolmogorov-Smirnov, 2,000 permutations). (i–k) Quantitative label-free proteomics workflow (n = 4 per group) (i), proteome volcano plot (j), and quantified glycogen-metabolizing enzymes in vitro and in vivo (k). (l and m) GAA protein induction in vivo in KP tumor-bearing lungs (n = 8 normoxic, 5 hypoxic tissue pieces from 3 and 2 mice; p = 0.02) (l) and in vitro in A549 cells (n = 4 per group; p = 1.0 × 10⁻⁵) (m). Data are mean ± SEM. (n) Model illustrating hypoxia-driven lysosomal GAA induction and subsequent glycogen catabolism supporting tumor growth. Statistics: pairwise comparisons were performed using two-tailed Student’s t tests with Benjamini-Hochberg correction; time course evaluated by permutation ANOVA. Proteomic fold changes were computed from per-sample normalized abundances. *p < 0.05, **p < 0.01, ***p < 0.001, ****p < 0.0001. See also Figures S6–S8.

Metabolic profiling revealed a distinct, time-dependent response to hypoxic stress. At 8 hours, total cellular glycogen increased significantly (1.80-fold), with accumulation evident across all measured chain lengths (Figures 4B and 4C). However, as the hypoxic exposure continued to 24 hours, this expanded glycogen pool was rapidly consumed (24-to-8-hour ratio 0.60), returning to a level statistically indistinguishable from baseline (1.09-fold; Figure 4C). Coinciding with this rapid glycogen depletion at 24 hours, upstream glycolytic and pentose-phosphate intermediates accumulated significantly, including fructose-1,6-bisphosphate (2.12-fold), phosphoenolpyruvate (9.24-fold), and 6-phosphogluconate (1.84-fold; Figure 4F). During this same 24-hour window, heavily utilized biosynthetic amino acids declined, including aspartate (0.45-fold), glutamine (0.60-fold), glutamate, and alanine (0.62-fold), alongside reduced glutathione (0.62-fold; Figures 4D and 4E; see Figure S6). Meanwhile, downstream pyruvate, lactate, and citrate pools remained highly stable throughout the exposure. Thus, hypoxic cancer cells actively synthesize glycogen during the early phases of stress and rapidly expend it as the hypoxia is sustained.

Furthermore, to provide additional evidence regarding the metabolic destination of this consumed glycogen carbon, we constructed a kinetic network model incorporating 35 nodes and 38 reactions (Methods). Forward simulations successfully reproduced the experimental metabolite pools (sum of squared errors 0.15), mapping the glycogen depletion into upper glycolysis and the pentose phosphate pathway (Figure 4G; see Figure S7). Metabolite-set enrichment independently supported this flux route, showing the activation of glycolysis (NES 1.50) and the pentose phosphate pathway, alongside the depletion of amino acid biosynthesis pathways (Figure 4H). Collectively, these in vitro data point toward hypoxic cancer cells dynamically mobilizing glycogen into anabolic pathways to fuel continued growth under metabolic stress, rather than merely storing it.

### Hypoxia induces the lysosomal enzyme GAA in cells and tumors

To explore the specific enzymatic route executing this glycogen mobilization, we performed quantitative proteomics on A549 cells cultured under normoxia and 8% oxygen (n = 4; Figures 4I and 4J). Among the nine quantified glycogen-metabolizing enzymes, lysosomal acid α-glucosidase (GAA) emerged as the most significantly induced target. Specifically, GAA increased by 1.92 log2 units (3.8-fold), vastly exceeding the induction of cytosolic glycogen phosphorylase (PYGL: 0.71 log2) and UGP2 (0.40 log2; Figure 4K). Concurrently, glycogen synthase (GYS1) decreased (−1.21 log2), while PGM1, PGM2, PYGB, GYG1, and GBE1 remained unchanged.

Interestingly, this GAA induction occurred without the concurrent activation of the canonical HIF-1α pathway. Under 8% oxygen, HIF-1α was not stabilized, and the canonical downstream target PDK3 remained unchanged, whereas CoCl_2_ treatment readily stabilized HIF-1α in these same cells (Figure S8). Consequently, GAA induction under moderate systemic hypoxia appears to uncouple from classical HIF transcription. By hydrolyzing glycogen within lysosomes, GAA yields free glucose that can efficiently feed upper glycolysis and the pentose phosphate pathway (Figure 4N)^32,33^.

To provide additional in vivo evidence, we performed quantitative proteomics on the tumor-bearing KP mouse lungs (n = 8/5). This confirmed a tumor-specific Gaa induction in vivo (1.77 log2 units; 3.4-fold increase; Figure 4L). Furthermore, Gaa was the only altered glycogen enzyme out of the ten quantified in vivo (Figure 4K), as Pygb, Gys1, Pygl, Pygm, Pgm1, Pgm2, Phkb, Ugp2, and Gbe1 were all unchanged. Collectively, both in vitro and in vivo models point toward the selective upregulation of lysosomal GAA under systemic hypoxia (Figures 4L and 4M).

### Tumor-specific deletion of GAA blunts the hypoxic response and eliminates the growth advantage

To evaluate whether this GAA induction is functionally required for hypoxia-driven tumor progression, we generated Kras^LSL-G12D/+^;Trp53^fl/fl^;Gaa^fl/fl^ (KP Gaa⁻/⁻) mice (Figure 5A)^29,30^. By utilizing adenoviral Cre delivery, we selectively excised Gaa solely within transformed lung epithelial cells, preserving normal host GAA expression and thereby avoiding broader systemic complications^32^. Immunofluorescence confirmed the robust presence of GAA signal in GAA-intact KP tumors and its subsequent absence within the KP Gaa⁻/⁻ tumors (Figures 5B and S9A). Furthermore, to verify that this genetic deletion effectively abolished substrate hydrolysis rather than merely eliminating antibody-detectable protein, we measured the enzymatic release of 4-methylumbelliferone (4-MU) from 4-methylumbelliferyl α-D-glucoside. This functional assay confirmed a significant reduction in GAA activity, dropping from approximately 53 nmol 4-MU/mg protein in GAA-intact KP tissue to approximately 23 nmol 4-MU/mg protein in KP Gaa⁻/⁻ tissue (n = 3 mice per group; Figure 5C). While this residual activity remained above the absolute baseline established by whole-body germline Gaa⁻/⁻ animals (approximately 3 nmol 4-MU/mg protein), it aligns with the expected retained activity from non-malignant cells that did not undergo Cre-mediated recombination. Notably, wild-type control tissue registered approximately 26 nmol 4-MU/mg protein.

**Figure 5.**
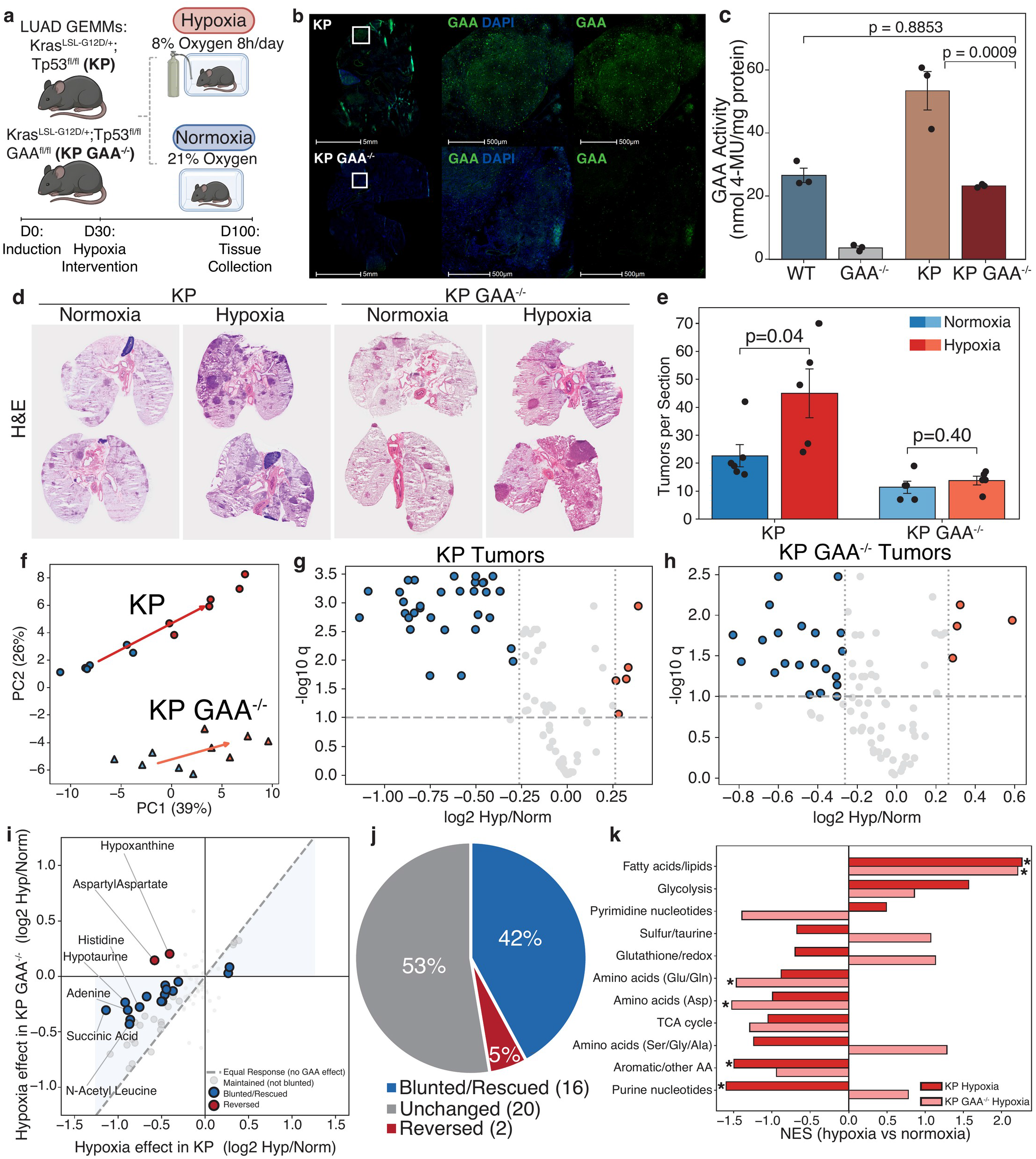
Tumor-specific deletion of GAA blunts the hypoxic response of KP lung tumors. (a) Experimental design for adenoviral Cre-mediated tumor induction and tumor-specific Gaa excision in Kras^LSL-G12D/+^;Trp53^fl/fl^ (KP) and Kras^LSL-G12D/+^;Trp53^fl/fl^;Gaa^fl/fl^ (KP Gaa⁻/⁻) mice. (b) Immunofluorescence of GAA (green) and DAPI (blue) in KP versus KP Gaa⁻/⁻ tumors (scale bars: overview 5 mm, inset 500 µm). (c) GAA enzyme activity measured via 4-methylumbelliferyl α-D-glucoside (4-MU) cleavage in wild-type, whole-body germline Gaa⁻/⁻, KP, and KP Gaa⁻/⁻ tissues (n = 3 mice per group). Bars represent group means with individual biological replicates overlaid. (d and e) Representative H&E histology (d) and quantification of tumors per section (e) under normoxia and hypoxia (n = 5–6 mice per group; KP p = 0.04, KP Gaa⁻/⁻ p = 0.40). Data are mean ± SEM with individual mice overlaid. (f) Principal component analysis of tumor spatial metabolomes following empirical Bayes batch correction (n = 5–6 tumors per group). Arrows indicate mean vector displacement from normoxia to hypoxia for each genotype. (g and h) Volcano plots mapping the hypoxic metabolic response in GAA-intact KP tumors (n = 38 shifted metabolites) (g) and KP Gaa⁻/⁻ tumors (n = 25 shifted metabolites) (h) (q < 0.1, ≥1.2-fold). (i) Hypoxia effect size in KP Gaa⁻/⁻ tumors plotted against the effect size in KP tumors for the 38 hypoxia-responsive metabolites. Dashed line denotes identity. (j) Classification of the 38 hypoxia-responsive metabolites in KP Gaa⁻/⁻ tumors (16 blunted/rescued, 20 maintained, 2 reversed). (k) Metabolite-set enrichment analysis of the hypoxic response within each genotype. Asterisks denote permutation p < 0.05. Statistics: slide batch effects were corrected within genotype using empirical Bayes. Genotype-by-hypoxia metabolite interactions were evaluated by ordinary least squares with Benjamini-Hochberg FDR correction. Across the metabolome, hypoxia displaced KP tumors 1.43-fold farther than KP Gaa⁻/⁻ tumors (p = 0.079). See also Figures S9 and S10.

Having established this functional deletion, we next assessed tumor burden. While systemic hypoxia successfully doubled tumor multiplicity in GAA-intact KP mice (1.99-fold increase; Figure 5E), this growth advantage was not observed in the KP Gaa⁻/⁻ cohort. Without tumor-cell-autonomous GAA, hypoxia did not significantly stimulate tumor growth (1.20-fold increase; Figures 5D and 5E). These data suggest that GAA is an important functional component of the hypoxia-mediated growth advantage.

To provide additional insight into the underlying metabolic impact of this genetic deletion, we mapped 87 metabolites across 22 tumors using spatial MALDI mass spectrometry imaging (GAA-intact KP: 6/6; KP Gaa⁻/⁻: 5/5). After correcting slide batch effects within each genotype using empirical Bayes (r = 0.992; Figures S9C–E), principal component analysis revealed a shortened hypoxic metabolic trajectory in the Gaa-deficient tumors (Figure 5F). In GAA-intact tumors, hypoxia induced broad shifts, significantly altering 38 metabolites, 33 of which decreased (Figure 5G). However, in KP Gaa⁻/⁻ tumors, the same exposure altered only 25 metabolites (Figure 5H). Across all 38 hypoxia-responsive metabolites, the median effect size in KP Gaa⁻/⁻ tumors dropped to just 0.51 of the intact response (Figure 5I; Figure S10). Consequently, tumor-specific GAA deletion reduced the overall magnitude of the hypoxic metabolic response.

Furthermore, classifying the 38 hypoxia-responsive metabolites by their behavior in KP Gaa⁻/⁻ tumors revealed 16 blunted or rescued, 20 maintained, and 2 reversed (Figure 5J). For instance, succinate depletion under hypoxia weakened from −1.14 log2 in GAA-intact KP tumors to −0.30 log2 in KP Gaa⁻/⁻ tumors. Similarly, hypotaurine shifted from −0.92 to −0.23, and hypoxanthine reversed entirely from a −0.41 depletion to a +0.20 accumulation. Pathway enrichment analysis highlighted the selective suppression of biosynthetic networks running downstream of free glucose (Figure 5K). Purine nucleotide synthesis, aromatic amino acids, serine-glycine-alanine metabolism, glutathione redox buffering, and glycolysis were all blunted or reversed in the KP Gaa⁻/⁻ tumors. Specifically, purine synthesis reversed from a normalized enrichment score (NES) of −1.60 in GAA-intact tumors to +0.78 in KP Gaa⁻/⁻ tumors. Meanwhile, mitochondrial pathways, including the tricarboxylic-acid cycle, aspartate synthesis, and glutamine-glutamate metabolism, remained intact or expanded in the absence of GAA. Collectively, these data point toward systemic hypoxia accelerating lung adenocarcinoma progression through GAA-dependent lysosomal glycogen catabolism.

## Discussion

Systemic hypoxemia is an intrinsic, compounding feature of advancing lung adenocarcinoma. As primary pulmonary tumors expand, they compromise functional alveolar surface area and obstruct airway patency, which inevitably lowers systemic oxygen delivery^2^. This resulting systemic oxygen deprivation establishes a systemic hypoxemic environment that may feed back onto the tumor to fuel further progression ^3^. Common clinical comorbidities, such as COPD and obstructive sleep apnea, actively compound this physiological burden and strongly correlate with reduced patient survival^6,7,10,11,12,13^. Utilizing a controlled in vivo model, our data suggest that systemic hypoxemia alone is biologically sufficient to increase lung tumor multiplicity. Furthermore, our metabolic profiles indicate that hypoxic tumors may sustain this rapid growth by dynamically mobilizing glycogen through lysosomal acid α-glucosidase (GAA).

Normal lung parenchyma and pulmonary tumors mount fundamentally different metabolic responses to systemic hypoxia. In non-malignant lung tissue, the metabolome remains highly stable, with glycogen representing the singular expanding carbon pool. When evaluating tumor-bearing lungs, bulk tissue homogenates often fail to resolve these specific metabolic shifts because the dense normal parenchyma dilutes the tumor signal^31^. Spatial mass spectrometry imaging elegantly addresses this by isolating the tumor compartment, revealing marked glycogen accumulation alongside coordinated, tumor-specific central carbon reprogramming^21,31^.

Cancer cells can degrade glycogen via two distinct enzymatic pathways. Cytosolic glycogen phosphorylase releases glucose-1-phosphate to sustain glycolysis during severe, acute anoxia^22,23,34,35^. In contrast, lysosomal GAA degrades glycogen to free glucose during sustained metabolic stress and glycophagy^24,25,32^. Under the moderate, sustained systemic hypoxia modeled here, lung tumors selectively induced GAA without altering the expression of cytosolic phosphorylase isoforms, and notably, this occurred independently of canonical HIF-1α stabilization. We postulate that lung tumors partition these glycogen degradation routes based on specific metabolic destinations. The cytosolic pathway rapidly provides glucose-1-phosphate to generate ATP, whereas the lysosomal pathway yields free glucose to preferentially support the pentose phosphate pathway, generating NADPH and fueling anabolic synthesis^33^. Consistent with this model, tumor-specific deletion of GAA selectively blunted purine nucleotide, serine, and glutathione pathways without broadly impairing the mitochondrial tricarboxylic-acid cycle.

These findings expand a rapidly evolving paradigm: glycogen functions as a highly adaptable, context-specific driver of lung adenocarcinoma. We previously established that lung cancer cells localize glycogen to the nucleus, where glycogen phosphorylase mobilizes it to directly modulate histone acetylation and epigenetic states^36^. Furthermore, recent work demonstrated that systemic nutritional excess, driven by a high-fat, high-carbohydrate Western diet, actively expands lung tumor glycogen reserves to accelerate progression^17^. The present study reveals that tumors similarly exploit systemic respiratory stress, but through a fundamentally distinct biochemical route. While dietary excess fuels general accumulation and nuclear stores regulate epigenetics, systemic hypoxia specifically triggers lysosomal glycophagy via GAA to sustain anabolic flux under metabolic crisis. Collectively, these paradigms provide additional evidence that lung tumors deploy compartmentalized glycogen metabolism to overcome diverse physiological challenges, shifting their degradation machinery to match the specific host-level stress.

Primary lung tumors generate systemic hypoxemia, and pre-existing respiratory comorbidities appear to amplify this burden to the detriment of patient survival. Resected human tumors from patients with sleep apnea exhibit elevated glycogen levels that closely align with our animal models. Collectively, these findings suggest a dual clinical vision: aggressively managing clinical hypoxemia through CPAP or supplemental oxygen could biologically starve the tumor of its hypoxic fuel, while directly inhibiting lysosomal GAA could represent an actionable therapeutic strategy to suppress lung adenocarcinoma progression in vulnerable, previously adapted patient populations.

### Limitations of the study

Our clinical cohorts were identified using diagnostic billing codes, which inherently lack direct physiological measurements of systemic oxygenation, apnea-hypopnea indices, or nocturnal desaturation. Consequently, granular variables such as precise smoking pack-years and specific cancer therapy regimens could not be fully controlled within these retrospective records. In the laboratory, our experimental design specifically isolated systemic hypoxemia using a controlled chamber to evaluate its singular biological contribution. However, progressive clinical comorbidities such as COPD and obstructive sleep apnea introduce complex, compounding variables, including chronic airway inflammation, hypercapnia, adrenergic stress, and rapid desaturation-reoxygenation cycles^37,38,39,40^. To build upon the current study, it will be necessary to establish the distinct impacts of these parallel factors on tumor progression and metabolism using empirical animal models that specifically recapitulate these conditions, such as cyclic intermittent hypoxia paradigms for sleep apnea. Furthermore, while our continuous 8% oxygen exposure established a robust baseline, future studies evaluating different oxygen percentages will help clarify how varying severities of hypoxemia modulate this metabolic response. Alongside this physiological complexity, human lung adenocarcinoma is highly mutationally heterogeneous. Because our mechanistic in vivo investigations utilized the canonical KP genetic model, future studies exploring whether GAA-dependent lysosomal glycophagy is conserved across other genomic subsets, such as STK11- or EGFR-mutant tumors, will help provide additional evidence for its broad translational potential. Finally, while spatial mass spectrometry imaging successfully localized these metabolic pools, future investigations incorporating in vivo ¹3C-glucose isotopic tracing will be beneficial to fully map the dynamic carbon flux trajectories feeding this systemic adaptation. Ultimately, translating these mechanistic insights into clinical application will require evaluating targeted therapeutic interventions. Future preclinical studies exploring the efficacy of hyperbaric oxygen therapy to physiologically counteract systemic hypoxemia, alongside the repurposing of pharmacological GAA inhibitors, currently utilized in the management of rare diseases, will help provide additional evidence for targeting this lysosomal vulnerability in lung adenocarcinoma.

## STAR★METHODS

### RESOURCE AVAILABILITY

#### Lead contact

Further information and requests for resources and reagents should be directed to and will be fulfilled by the lead contact, Ramon C. Sun.

#### Materials availability

The KP;Gaa^fl/fl^ (KP Gaa^−/−^) mouse line generated by crossing the KP model to the Gaa^fl/fl^ line is available from the lead contact under a standard material transfer agreement. This study did not generate other new unique reagents.

#### Data and code availability

All raw data reported in this paper, including the MALDI mass spectrometry imaging data and the label-free proteomics data, together with the processed metabolite and glycogen chain-length matrices and the proteomics abundance tables, will be publicly available at sunlabresource.com upon publication. Patient-level clinical data from the OneFlorida+ Clinical Research Consortium are not publicly available because of patient privacy protections and are accessible through application to the consortium. This paper does not report original code; all analyses utilized the software and published algorithms described in the Method Details and Quantification and Statistical Analysis sections.

### EXPERIMENTAL MODEL AND STUDY PARTICIPANT DETAILS

#### Mice

All animal experiments were approved by the Institutional Animal Care and Use Committee at the University of Florida (protocol 202200000686) and were conducted in accordance with institutional ethical guidelines. Mice were housed in a climate-controlled room. Mice were 15 weeks of age (approximately 105 days) at tumor induction, which defined experimental day 0; the 30-day tumor-formation and 70-day hypoxia-exposure periods followed this day-0 induction. The KP model (Kras^LSL-G12D/+^; Trp53^fl/fl^) was crossed with the Gaa^fl/fl^ line from the Fuller laboratory to generate the KP Gaa^−/−^ model (Kras^LSL-G12D/+^; Trp53^fl/fl^; Gaa^fl/fl^). Both male and female mice were included because sex differences have not been reported in these models. Mice were randomly assigned to treatment groups, and investigators were double-blinded until statistical evaluation was complete.

#### Cell lines

A549 human lung adenocarcinoma cells (ATCC CCL-185) were maintained in 10 mM glucose Dulbecco’s modified Eagle medium (DMEM) supplemented with 10% fetal bovine serum (Thermo Fisher Scientific) and 1% penicillin-streptomycin (Thermo Fisher Scientific) at 37 °C in a humidified 5% CO₂ incubator. All experiments used low-passage cells (≤ p10) from the original ATCC stock, passaged at approximately 80% confluence with 0.25% trypsin-EDTA. Cells were confirmed mycoplasma-negative every 6 months. For every experiment, cells were seeded the day before treatment and allowed to adhere overnight so that each experiment began on a fully adhered monolayer; seeding format and density were matched to the downstream assay.

#### Human participants: MALDI tissue cohort

Formalin-fixed human non-small cell lung cancer tissue blocks from patients with and without a documented clinical history of sleep apnea (n = 5 per group) were analyzed by MALDI mass spectrometry imaging. Groups were matched on stage and grade. All human tissue samples were deidentified, and the study was determined to be IRB exempt. The full clinicopathologic characteristics of the 10 cases are provided in Table S1.

#### Human participants: OneFlorida clinical cohort

A retrospective cohort of patients with incident lung cancer was assembled from the OneFlorida electronic health record network linked to tumor registry data. Two pre-existing respiratory exposures were evaluated in parallel within this cohort, obstructive sleep apnea and chronic obstructive pulmonary disease, each analyzed as a separate propensity-score-matched comparison against its own matched controls. The analysis used deidentified records and was determined to be IRB exempt. Cohort construction, exposure definitions, and statistical analysis are detailed under Method Details and Quantification and Statistical Analysis.

### METHOD DETAILS

#### AdenoCre induction

Adeno-Cre adenoviral vectors were obtained from the Viral Vector Core at the University of Iowa (Iowa City, IA, USA); vector concentration and functionality were confirmed by the manufacturer using quantitative PCR and plaque assays. For anesthesia, mice received a ketamine-xylazine-saline mixture at 10 µL g^−1^. Anesthetized mice were placed supine on a heating pad to maintain core body temperature. Adeno-Cre viral solution was delivered intranasally by pipette in a total volume of 62.5 µL (61.8 µL Advanced DMEM, 0.3 µL CaCl₂, and 0.416 µL Adeno-Cre) at a titer of 1 × 10^10^ plaque-forming units per milliliter. Mice were then monitored daily for signs of distress, body-weight change, and general health, in compliance with IACUC guidelines, until endpoint euthanasia or 100 days post-induction.

#### Systemic hypoxia exposure

Mice were induced with Adeno-Cre and tumors were allowed to form for 30 days. Half of the animals were then exposed to 21% oxygen (defined as normoxia) and half to 8% oxygen (defined as hypoxia). Hypoxia was delivered using an OxyCycler A41OV atmosphere-controlled chamber and software. Each hypoxic bout lasted 8 h and was applied 5 days per week for 70 days, until the tumor endpoint was reached.

#### Whole-body plethysmography

Whole-body plethysmography (WBP; DSI, Buxco FinePointe) was used to measure respiratory rate, tidal volume, and minute ventilation. Before day 0, mice were acclimated to the chambers for 30 min on two consecutive days. On day 0, mice received a 30 min acclimation, a 15 min baseline normoxic recording (21% oxygen), a 10 min hypoxic-hypercapnic exposure (8% oxygen, 5% CO₂), and a 10 min post-exposure normoxic recording. On days 1, 7, and 14, mice received a 30 min acclimation, a 15 min baseline hypoxic recording (8% oxygen), a 10 min hypoxic-hypercapnic exposure (8% oxygen, 5% CO₂), and a 10 min re-acclimation to hypoxia. Normoxic mice followed the same protocol with normoxic rather than hypoxic exposure.

#### Immunofluorescence

Fresh-frozen lung tissues were fixed in 1% paraformaldehyde. Antigen retrieval was performed at 95 °C in an alkaline buffer for 30 min. Tissues were stained with a primary antibody against acid α-glucosidase (GAA; Abcam ab102815) and detected with a fluorescent goat anti-rabbit IgG secondary. Slides were scanned at 40× on a VS200 slide scanner (Olympus) and reviewed in HALO software (Indica Labs). GAA immunofluorescence was assessed qualitatively for tumor-cell presence versus absence in KP and KP Gaa^−/−^ tumors (Figures 5 and S9).

#### Cell hypoxia exposure

Normoxic control cells were held in a standard 5% CO₂ incubator at ambient (21%) oxygen. Hypoxic cells were held in a BioSpherix hypoxia chamber (Xvivo System X3) at 8% O₂, 5% CO₂, 37 °C, and 95% humidity. An oxygen concentration of 8% was chosen to match the in vivo hypoxia experiment. Cells were exposed for 8 or 24 h as indicated and harvested simultaneously on ice at the end of exposure to limit reoxygenation. For the western-blot positive control, cobalt chloride (CoCl₂) was applied at 400 µM in complete medium for 24 h to stabilize HIF-1α. Cells for western blot were plated in 6-well plates at 300,000 cells per well (Nest Scientific) with n = 3 replicates per condition; proteomics cells were plated in 10 cm dishes with n = 4 replicates per condition; spatial-metabolomics cells were seeded on eight-chamber well slides with n = 7 or 8 wells per condition.

#### Protein extraction and BCA quantification

At the end of treatment, each plate was placed on ice, washed with ice-cold phosphate-buffered saline (PBS), and lysed in RIPA buffer with HALT protease-phosphatase inhibitors (Thermo Fisher Scientific). Lysates were centrifuged at 15,000g for 15 min at 4 °C and supernatants retained. Total protein was quantified by bicinchoninic acid assay (Pierce BCA Protein Assay, Thermo Fisher Scientific) against a bovine serum albumin standard curve (0–2,000 µg/mL, run in duplicate). Lysates were assayed at a 1:10 dilution, absorbance was read at 562 nm, and concentrations were interpolated from a linear regression of the blank-subtracted standards (R^2^ = 0.985) and corrected for the dilution factor, giving lysate concentrations of approximately 2.7–3.5 mg/mL.

#### GAA enzyme activity assay (4-methylumbelliferyl α-D-glucoside)

Tissue was cryosectioned and collected into a microcentrifuge tube and protein extraction buffer was added. After one hour, tubes were centrifuged and the supernatant was retained. Protein concentration was determined with the Pierce BCA Protein Assay (Thermo Fisher Scientific) and samples were normalized to 1 mg/mL. In a 96-well plate, 10 µL of either 4-methylumbelliferone standard or protein sample was combined with a mixture of MilliQ water, sodium acetate buffer, and 4-methylumbelliferyl α-D-glucoside. Plates were incubated for 1 h at 37 °C, the reaction was stopped with sodium carbonate buffer, and fluorescence was measured using a fluorescence microplate reader.

#### Jess capillary western immunoassay

Target proteins were resolved and detected by automated capillary immunoassay on a Jess system (ProteinSimple, Bio-Techne) using the 12–230 kDa separation module with chemiluminescent detection. For each sample, lysate was diluted to 1.56 mg/mL in 0.1× Sample Buffer and combined 4:1 (v/v) with 5× fluorescent master mix; 1× Sample Buffer and 40 mM dithiothreitol (final) were then added, giving a denatured working concentration of 1.25 mg/mL. Samples were denatured at 95 °C for 5 min in a thermal cycler with a heated lid and centrifuged at 10,000g for 2 min at room temperature, and the supernatant was loaded. Plates were prepared per the manufacturer’s protocol: 5 µL of each denatured sample (1.25 mg/mL) was dispensed per capillary well, followed by Antibody Diluent 2, primary antibody, secondary antibody or streptavidin-HRP for the ladder, and luminol-peroxide. Separation, immobilization, immunoprobing, and signal acquisition were run under default 12–230 kDa parameters and quantified in Compass for Simple Western using Gaussian-fit peak areas. Each target was multiplexed within the same capillary with β-actin as the loading control by co-incubating target and β-actin primary antibodies (both raised in rabbit) and resolving them as separate bands. Primary antibodies were anti-HIF-1α (rabbit, Cell Signaling Technology 36169, 1:50) and anti-β-actin (rabbit, GeneTex GTX637675, 1:1000); the secondary was anti-rabbit-HRP (ProteinSimple, ready-to-use).

#### MALDI mass spectrometry imaging: tissue small molecules

Lung tissue was surgically removed and slow-frozen in liquid nitrogen. Cryosections (10 µm) were prepared from fresh-frozen lungs and mounted directly onto microscope slides without optimal-cutting-temperature compound. Sections were promptly dried in a desiccator to halt metabolic activity. N-(1-naphthyl)ethylenediamine dihydrochloride matrix (7 mg/mL in 70% methanol) was applied with an HTX M5 sprayer. Imaging was performed on a Bruker timsTOF fleX (laser 10,000 Hz, 60% energy, 300 shots per pixel, 50 µm raster and spot size), targeting small molecules and lipids. Spectra were acquired in negative-ion mode across m/z 50–2,000.

#### MALDI mass spectrometry imaging: glycogen, fresh-frozen tissue

Fresh-frozen cryosections (10 µm) were prepared and desiccated as above, then fixed in 10% neutral buffered formalin. Antigen retrieval was performed in citraconic anhydride buffer (pH 3) using a vegetable steamer for 30 min. After cooling, the buffer was incrementally exchanged with water and slides were desiccated before enzymatic digestion. Isoamylase (3 units per slide) was applied with the HTX spray station, and slides were incubated at 37 °C for 2 h before matrix application. Imaging was performed on a Bruker timsTOF fleX (laser 10,000 Hz, 60% energy, 300 shots per pixel, 50 µm raster and spot size), targeting glycogen. Spectra were acquired in positive-ion mode across m/z 600–4,000.

#### MALDI mass spectrometry imaging: glycogen, FFPE tissue

The human lung cancer tissue cohort (Figures 3O–Q) was formalin-fixed and paraffin-embedded. FFPE sections (4 µm) were mounted for MALDI imaging following established protocols. Slides were heated at 60 °C for 1 h, deparaffinized in xylene, and rehydrated through graded ethanol into water. Antigen retrieval, isoamylase digestion, and matrix application were performed as for fresh-frozen glycogen imaging. Imaging was performed on a Bruker timsTOF platform (laser 1,000 Hz, intensity 200 a.u., 50 µm spot size) across m/z 600–4,000. Data were processed in Bruker SCiLS Lab.

#### MALDI mass spectrometry imaging: cultured cells

Spatial metabolomics of cultured cells was performed as previously described (Clarke et al., 2025). A549 cells were seeded at 35,000 cells per well on eight-chamber well slides (Lab-Tek), allowed to adhere and grow overnight, and exposed to normoxia (21% O₂) or 8% O₂ for 8 or 24 h. At endpoint, medium was aspirated, cells were washed with 0.1× PBS to remove salts, and metabolism was quenched by dehydration in a vacuum desiccator, where slides were held until matrix application the following day. Metabolome and glycogen were imaged sequentially from the same wells. For small molecules, slides were coated with N-(1-naphthyl)ethylenediamine dihydrochloride (NEDC) matrix (7 mg/mL in 70% methanol) using an HTX M5 sprayer and imaged on a Bruker timsTOF fleX (10,000 Hz, 60% energy, 1,000 shots per pixel, 100 µm pixel size) across m/z 50–1,000 in negative-ion mode. NEDC matrix was then removed with ice-cold 100% methanol, slides were processed for glycogen as above, α-cyano-4-hydroxycinnamic acid (CHCA) matrix was applied, and glycogen was imaged on the timsTOF fleX in positive-ion mode. Images were processed in SCiLS Lab (Bruker); steady-state small-molecule and lipid features were exported through the SCiLS API with total-ion-current (TIC) normalization, and the glycogen degree-of-polymerization ladder was exported as raw peak areas.

#### Label-free proteomics: sample preparation

A549 cells (control and 8% O₂ for 8 h; n = 4 per condition) and mouse lung tissue (normoxia n = 8 and hypoxia n = 5) were processed for bottom-up proteomics. For A549, medium was preconditioned at 8% oxygen for 6 h before treatment; cells were harvested, washed in PBS, pelleted (500g, 5 min), and snap-frozen in liquid nitrogen. Proteins were extracted and digested using the EasyPep MS Sample Prep Kit (Thermo Fisher Scientific). Total protein concentration was determined on a Qubit fluorometer (Invitrogen), and an equal amount of total protein per sample was taken for digestion. Samples were reduced with 0.1 M dithiothreitol in 100 mM ammonium bicarbonate at 56 °C for 30 min, alkylated with 55 mM iodoacetamide in 100 mM ammonium bicarbonate at room temperature in the dark for 30 min, and digested with sequencing-grade trypsin/Lys-C (Promega) at 70 °C for 1 h. Digestion was quenched with 0.5% trifluoroacetic acid, and tryptic peptides were analyzed by LC-MS/MS immediately.

#### Nano-LC-MS/MS acquisition

Nano-liquid chromatography tandem mass spectrometry was performed on a Thermo Scientific Q Exactive HF Orbitrap mass spectrometer equipped with an EASY-Spray nanospray source and operated in positive-ion mode, coupled to an UltiMate 3000 RSLCnano system. Mobile phase A was water with 0.1% formic acid and mobile phase B was acetonitrile with 0.1% formic acid; the loading-pump mobile phase was water with 0.1% trifluoroacetic acid. For each sample, 5 µL was injected onto a µPAC C18 trapping column at 10 µL/min, held for 3 min, and washed with 1% B to desalt and concentrate the peptides. Peptides were eluted onto a 110-cm µPAC analytical column maintained at 40 °C at 750 nL/min for the first 15 min and then 250 nL/min, using a gradient of 1% to 20% B over 100 min and then to 45% B over 20 min, for a total run time of 150 min. The EASY-Spray source operated at 1.5 kV with a capillary temperature of 200 °C. MS/MS was acquired with a data-dependent method: a full MS1 scan from m/z 375 to 1,575 at 60,000 resolution followed by MS2 scans at 15,000 resolution on the 15 most abundant precursors, with HCD fragmentation at a stepped normalized collision energy of 28 and a 4 m/z isolation window; singly charged ions were excluded, dynamic exclusion was enabled, and m/z 445.12003 served as an internal lock mass.

#### Label-free proteomics: database search and quantification

All MS/MS spectra were searched with the CHIMERYS node (MSAID) in Proteome Discoverer 3.2.0.450 (Thermo Fisher Scientific) using the CWF_Comprehensive_Enhanced Annotation_LFQ_and_Precursor_Quan workflow. Spectra were searched against the UniProt Homo sapiens reference proteome (sp_incl_isoforms, TaxID 9606, release 407; A549) or the Mus musculus reference proteome (sp_incl_isoforms, TaxID 10088 and sub-taxonomies, release 406; mouse lung), each concatenated with a universal contaminants FASTA, with trypsin as the digestion enzyme, carbamidomethylation of cysteine as a static modification, and oxidation of methionine as a dynamic modification. PSM and peptide identifications were validated at a strict 1% FDR by q-value filtering, and high-confidence peptides (minimum length 6) were retained. Protein grouping used strict parsimony. Precursor-ion-intensity label-free quantification was performed in Proteome Discoverer using unique and razor peptides, with abundances normalized to total peptide amount per sample without additional scaling. Gene Ontology and pathway annotations were added via the Thermo Protein Annotation server, and results were filtered to master proteins at high FDR confidence. Downstream fold changes were recomputed from the per-sample normalized abundances rather than from the Proteome Discoverer ratio column (see Quantification and Statistical Analysis).

#### OneFlorida retrospective cohort: design and exposure

Patients with incident lung cancer were identified using OneFlorida electronic health records linked to tumor registry data. Incident lung cancer was defined as the first tumor-registry record carrying an ICD-O site code of C34* or a numeric site code of 340 to 349. The index date (time zero) was the date of lung cancer diagnosis. Eligibility required at least 365 days of observable clinical history before diagnosis to support baseline covariate ascertainment, and, for the COPD analysis, nonnegative post-diagnosis follow-up. Patients were followed from diagnosis until death, disenrollment, or administrative censoring on 1 January 2024, and the primary endpoint was all-cause mortality. Pre-existing obstructive sleep apnea (OSA) was defined by a high-specificity ICD-based algorithm requiring at least two OSA-consistent diagnosis codes strictly before time zero; patients not meeting this definition were classified as unexposed. Baseline covariates for the OSA analysis, defined using only pre-diagnosis information, comprised age at diagnosis, sex, race and ethnicity, cancer stage, baseline healthcare utilization (encounter and hospitalization counts), and the Charlson Comorbidity Index. Pre-existing chronic obstructive pulmonary disease (COPD) was defined in parallel by a high-specificity ICD-based algorithm requiring at least two qualifying COPD diagnosis events strictly before time zero, with qualifying ICD-10-CM codes J44.0, J44.1, J44.9, and J44.89. Baseline covariates for the COPD analysis, likewise restricted to pre-diagnosis information, comprised age at diagnosis, sex, Hispanic ethnicity, tumor stage, congestive heart failure, chronic kidney disease, diabetes, obesity, and baseline healthcare utilization measured as encounter and hospitalization counts. The two exposures were ascertained using identical algorithmic criteria but matched on distinct covariate sets, and were analyzed as independent comparisons.

### QUANTIFICATION AND STATISTICAL ANALYSIS

#### Computational environment and reproducibility

Unless otherwise noted, imaging, proteomics, and modeling analyses were performed in Python (numpy, pandas, matplotlib) or GraphPad Prism. Group comparisons utilized a validated pooled-variance two-tailed Student’s t test, with multiple comparisons controlled by the Benjamini-Hochberg false-discovery-rate procedure (reported as q). Two significance thresholds were applied and are stated explicitly wherever they are used. For MALDI analyses, a biological effect was defined by a fold-change threshold of 1.2× with q < 0.05 for the primary group contrasts, whereas the discovery-level analyses reported here, namely volcano coloring for differential metabolite testing, temporal-trend clustering, and proteomic differential expression, applied a more permissive threshold of q < 0.1 as specified in the corresponding sections. Data are presented as mean ± SEM, with significance markers denoting *p or q < 0.05, **< 0.01, ***< 0.001, and ****< 0.0001. Animal-level and column analyses utilized two-tailed Student’s t tests or one- or two-way ANOVA as appropriate, with the specific test and replicate number detailed in the respective figure legends.

#### MALDI-MSI preprocessing and normalization

Image acquisition and primary spectral processing were performed in flexImaging v6.0 and SCiLS Lab v2024b (Bruker Daltonics). SCiLS Lab exports were parsed by removing the matrix peak and exact-duplicate feature rows, and features detected in at least 50% of regions in at least one group were retained. Isobaric features sharing an m/z were flagged as mass-ambiguous. Small-molecule and lipid intensities were total-ion-current (TIC) normalized and log10-transformed; where noted, Pareto scaling was additionally applied for multivariate and heatmap visualization. To account for spatial pseudoreplication, pixels were aggregated to a per-piece or per-tissue mean for each tissue comparison, and statistical significance was computed at the replicate level. The wild-type normal-lung arm included n = 5 mice per group. The KP whole tumor-bearing lung utilized the tissue piece as the analytical unit (n = 9 normoxic and 6 hypoxic pieces, from 3 and 2 mice, respectively), with heart pixels excluded. The tumor-compartment analysis included 5 large tumors per group, each from a different animal.

#### Differential metabolite testing and volcano analysis

Per-feature fold change was computed as the ratio of group means on per-replicate values. Significance was assessed using a pooled-variance Student’s t test on log10 values with Benjamini-Hochberg FDR across all detected features. Volcano plots display −log10(q) against log2 fold change, with features colored by direction at q < 0.1 and absolute fold change > 1.2×. Testing covered 71 features (70 small molecules plus total glycogen) in wild-type normal lung and 96 small molecules in the KP whole-lung and tumor-compartment datasets. A549 comparisons were conducted at the region level (n = 8 regions per group) at 8 h and 24 h of 8% O₂ versus normoxia.

#### Dimensionality reduction

Principal-component analysis was computed on log10 (and, where noted, Pareto-scaled) feature matrices. Pixel-level UMAP embeddings were computed on balanced pixel subsamples (2,500 pixels per group) after a sparse-pixel quality filter (at least 10 detected ions per pixel), and global mixing was quantified by the cross-group fraction among each pixel’s k nearest neighbors, where a value near 0.5 indicates full intermixing and no global metabolic separation. Canonical UMAP figures were generated with the umap-learn package.

#### Multivariate discriminant analysis

For the A549 steady-state metabolome (Figure S6), partial least-squares discriminant analysis (PLS-DA) was fit for normoxia versus pooled hypoxia on the log10 feature matrix, and class separation was summarized by RZY and QZ from cross-validation. Model significance was assessed by permutation with 300 label permutations, and variable importance in projection (VIP) scores identified the metabolites driving separation at VIP > 1.

#### Glycogen chain-length quantification

Glycogen was measured as a hexose degree-of-polymerization ladder (DP4 to DP21, spaced by 162 Da per glucosyl unit; DP7 at m/z 1,175), exported as raw peak areas without normalization. Total glycogen was the sum across the detected chain-length channels (17 channels were detected in normal lung and 18 in the tumor and A549 datasets). Chain-length distributions were compared both as absolute per-DP peak areas and as within-sample-normalized profiles (each sample scaled to its own total). For the A549 glycogen time course (2 oxygen levels × 3 timepoints, n = 7–8 regions per group), group differences were assessed by six-group permutation analysis of variance across both oxygen levels.

#### Batch correction and the genotype-by-hypoxia interaction (KP and KP Gaa^−/−^ cohort)

The KP versus KP Gaa^−/−^ interaction cohort consisted of 22 tumors (KP: 6 normoxic, 6 hypoxic; KP Gaa^−/−^: 5 normoxic, 5 hypoxic). Pixels were TIC-normalized, aggregated to per-tissue means, log10-transformed, and corrected for slide batch effects using a parametric empirical-Bayes model applied within each genotype separately, protecting condition as a covariate. The hypoxia effect within each genotype was tested by Student’s t test on the corrected log10 values with Benjamini-Hochberg FDR. The genotype-by-hypoxia interaction was evaluated per metabolite using ordinary least squares regression on the corrected data with a genotype-by-condition interaction term and Benjamini-Hochberg FDR. Among the KP hypoxia-responsive metabolites, responses were classified as reversed, blunted or rescued (magnitude <50% of KP), maintained, or amplified.

#### Metabolite-set and gene-set enrichment

Pathway enrichment utilized a weighted Kolmogorov-Smirnov running-sum enrichment score on features ranked by their signed Student’s t statistic for the relevant contrast. Significance was assessed by competitive permutation against 2,000 random feature sets of matched size to derive the normalized enrichment score (NES) and permutation p value, followed by Benjamini-Hochberg FDR correction. Metabolite sets included curated central-carbon, amino-acid, nucleotide, redox, and lipid groups drawn from detected features, retaining sets with at least three members. The glycogen set comprised the 18 chain-length channels computed on raw intensity; small-molecule sets were TIC-normalized. This procedure was similarly applied to protein-level enzyme and Gene Ontology gene sets for proteomics datasets.

#### Temporal-trend clustering

For the A549 hypoxia time course (Figure S7), metabolites with a significant temporal effect (one-way ANOVA q < 0.1) were grouped into temporal-trend clusters by k-means clustering of their z-scored time trajectories, and total glycogen was overlaid onto the resulting clusters to show that it co-clusters with metabolites that peak at 8 h and then decline.

#### Kinetic modeling of A549 glycogen utilization

To evaluate glycogen mobilization consistency with measured downstream pools, 8 h and 24 h A549 metabolome and glycogen data were integrated in a tracer-free pool-dynamics analysis. Ordinary-differential-equation network models, ranging from a reduced 5-pool model to a 35-node, 38-reaction central-carbon network with vectorized stoichiometry, were fit to the relative 8 h to 24 h pools by multi-start coordinate descent to model the measured node trajectories.

#### Proteomics quantification

Fold changes were recomputed from the per-sample normalized abundances after excluding contaminant and non-target-species entries, rather than from the Proteome Discoverer precomputed abundance-ratio column. Differential expression used log2 abundances with the Student’s t test and Benjamini-Hochberg FDR, and proteomic volcano plots colored proteins at |log2 fold change| > 0.58 (1.5×) and q < 0.1. Proteomics used n = 4 per group for A549 cells and n = 8 normoxic and 5 hypoxic samples for KP tumor-bearing lung. Protein-level enzyme-set and Gene Ontology GSEA used the same weighted-Kolmogorov-Smirnov, signed-t, 2,000-permutation procedure as the metabolite enrichment.

#### Human tissue glycogen imaging

Human FFPE lung cancer tissue glycogen images (control versus sleep apnea, n = 5 patients per group) were analyzed without intensity normalization to preserve the total-glycogen signal. Pixels were pooled per patient for patient-level summaries.

#### OneFlorida clinical cohort: propensity-score matching and survival analysis

A propensity score for pre-existing OSA was estimated by regularized logistic regression on all baseline covariates in the eligible cohort of 3,044 patients. Patients were matched 1:1 without replacement by nearest-neighbor matching on the logit of the propensity score using a caliper of 0.15 standard deviations of the logit propensity score, with additional exact matching on Stage IV status. The procedure yielded a matched cohort of 470 patients (235 with and 235 without pre-existing OSA), in which stage was identical by construction and sex, race, and ethnicity were well balanced. Survival time was measured from lung cancer diagnosis and summarized with Kaplan-Meier curves compared by the log-rank test, with hazard ratios from a Cox proportional-hazards model; both unadjusted and covariate-adjusted models were fit within the matched cohort to address residual differences in age and comorbidity.

A propensity score for pre-existing COPD was estimated in parallel by ridge logistic regression on the COPD baseline covariates in an eligible cohort of 3,028 patients, of whom 601 carried a COPD diagnosis before lung cancer. Patients were matched 1:1 without replacement by nearest-neighbor matching on the logit of the propensity score using a caliper of 0.15 standard deviations of the logit propensity score, with additional exact matching on collapsed tumor stage. The procedure yielded a matched cohort of 1,196 patients, 598 with and 598 without pre-existing COPD. Survival was analyzed as for the OSA cohort, with sex-stratified analyses and sensitivity analyses using crude, covariate-adjusted, and inverse-probability-of-treatment-weighted Cox regression.

#### Whole-body plethysmography analysis

Respiratory frequency, tidal volume, and minute ventilation were computed by the FinePointe software. Minute ventilation was normalized to body weight, and each animal contributed a single 5-min-average value per recording block and timepoint (n = 8 mice per arm). Groups were compared at each timepoint against baseline with data shown as mean ± SEM and exact p values reported.

## Acknowledgments

This study was supported by National Institutes of Health (NIH) grants R01CA266004, R01CA288696, R01AG066653, R01AG078702, and RM1NS133593 to R.C.S., and R35NS116824 to M.S.G. Additional support was provided by the Center for Advanced Spatial Biomolecule Research (CASBR) at the University of Florida College of Medicine. Large language models (Claude) were used for grammar checking and proofreading.

## Author Contributions

Conceptualization, H.A.C., C.J.S., H.P., Y.G., M.S.G., and R.C.S.; Methodology, R.C.S., M.S.G., C.W.V.K., H.A.C., T.R.H., K.B.B., D.D.F., and Y.G.; Formal Analysis, H.A.C., C.J.S., H.P., R.A.R., and Y.G.; Investigation, H.A.C., C.J.S., H.P., T.R.H., D.D.K., O.J., A.V., F.B., C.M.S., R.L., A.M.R., R.A.R., L.W., J.R., Y.W., and K.B.B.; Pathology Review, D.B.A.; Resources, C.W.V.K., K.B.B., D.B.A., M.C., D.D.F., B.J.B., M.S.G., and R.C.S.; Writing – Original Draft, R.C.S. and H.A.C.; Writing – Review & Editing, R.C.S., M.S.G., Y.G., C.W.V.K., K.B.B., D.B.A., M.C., D.D.F., and B.J.B.; Funding Acquisition, R.C.S., M.S.G., and Y.G.; Supervision, R.C.S., M.S.G., and Y.G.

## References

1. Bray, F., Laversanne, M., Sung, H., Ferlay, J., Siegel, R.L., Soerjomataram, I., and Jemal, A. (2024). Global cancer statistics 2022: GLOBOCAN estimates of incidence and mortality worldwide for 36 cancers in 185 countries. CA Cancer J. Clin. 74, 229–263.

2. Ernst, A., Feller-Kopman, D., Becker, H.D., and Mehta, A.C. (2004). Central airway obstruction. Am. J. Respir. Crit. Care Med. 169, 1278–1297.

3. Muz, B., de la Puente, P., Azab, F., and Azab, A.K. (2015). The role of hypoxia in cancer progression, angiogenesis, metastasis, and resistance to therapy. Hypoxia 3, 83–92.

4. Benjafield, A.V., Ayas, N.T., Eastwood, P.R., Heinzer, R., Ip, M.S.M., Morrell, M.J., Nunez, C.M., Patel, S.R., Penzel, T., Pépin, J.L., et al. (2019). Estimation of the global prevalence and burden of obstructive sleep apnoea: a literature-based analysis. Lancet Respir. Med. 7, 687–698.

5. GBD Chronic Respiratory Disease Collaborators (2020). Prevalence and attributable health burden of chronic respiratory diseases, 1990-2017: a systematic analysis for the Global Burden of Disease Study 2017. Lancet Respir. Med. 8, 585–596.

6. Park, H.Y., Kang, D., Shin, S.H., Yoo, K.H., Rhee, C.K., Suh, G.Y., Kim, H., Shim, Y.M., Guallar, E., Cho, J., et al. (2020). Chronic obstructive pulmonary disease and lung cancer incidence in never smokers: a cohort study. Thorax 75, 506–509.

7. Cheong, A.J.Y., Tan, B.K.J., Teo, Y.H., Tan, N.K.W., Yap, D.W.T., Sia, C.H., Ong, T.H., Leow, L.C., See, A., and Toh, S.T. (2022). Obstructive sleep apnea and lung cancer: a systematic review and meta-analysis. Ann. Am. Thorac. Soc. 19, 469–475.

8. Rodríguez-Roisin, R., Drakulovic, M., Rodríguez, D.A., Roca, J., Barberà, J.A., and Wagner, P.D. (2009). Ventilation-perfusion imbalance and chronic obstructive pulmonary disease staging severity. J. Appl. Physiol. 106, 1902–1908.

9. Jordan, A.S., McSharry, D.G., and Malhotra, A. (2013). Adult obstructive sleep apnoea. Lancet 383, 736–747.

10. McNicholas, W.T. (2017). COPD-OSA overlap syndrome: evolving evidence regarding epidemiology, clinical consequences, and management. Chest 152, 1318–1326.

11. Nieto, F.J., Peppard, P.E., Young, T., Finn, L., Hla, K.M., and Farré, R. (2012). Sleep-disordered breathing and cancer mortality: results from the Wisconsin Sleep Cohort Study. Am. J. Respir. Crit. Care Med. 186, 190–194.

12. Martínez-García, M.A., Campos-Rodriguez, F., Durán-Cantolla, J., de la Peña, M., Masdeu, M.J., González, M., Del Campo, F., Serra, P.C., Valero-Sánchez, I., Ferrer, M.J., et al. (2014). Obstructive sleep apnea is associated with cancer mortality in younger patients. Sleep Med. 15, 742–748.

13. Campos-Rodriguez, F., Martinez-Garcia, M.A., Martinez, M., Duran-Cantolla, J., de la Peña, M.L., Masdeu, M.J., Gonzalez, M., del Campo, F., Gallego, I., Marin, J.M., et al. (2013). Association between obstructive sleep apnea and cancer incidence in a large multicenter Spanish cohort. Am. J. Respir. Crit. Care Med. 187, 99–105.

14. Hensley, C.T., Faubert, B., Yuan, Q., Lev-Cohain, N., Jin, E., Kim, J., Jiang, L., Ko, B., Skelton, R., Loudat, L., et al. (2016). Metabolic heterogeneity in human lung tumors. Cell 164, 681–694.

15. Davidson, S.M., Papagiannakopoulos, T., Olenchock, B.A., Heyman, J.E., Keibler, M.A., Luengo, A., Bauer, M.R., Jha, A.K., O’Brien, J.P., Pierce, K.A., et al. (2016). Environment impacts the metabolic dependencies of Ras-driven non-small cell lung cancer. Cell Metab. 23, 517–528.

16. Mazumdar, J., Hickey, M.M., Pant, D.K., Durham, A.C., Sweet-Cordero, A., Vachani, A., Jacks, T., Chodosh, L.A., Kissil, J.L., Simon, M.C., et al. (2010). HIF-2alpha deletion promotes Kras-driven lung tumor development. Proc. Natl. Acad. Sci. USA 107, 14182–14187.

17. Clarke, H.A., Hawkinson, T.R., Shedlock, C.J., Medina, T., Ribas, R.A., Wu, L., Liu, Z., Ma, X., Xia, Y., Huang, Y., et al. (2025). Glycogen drives tumour initiation and progression in lung adenocarcinoma. Nat. Metab. 7, 952–965.

18. Liu, Q., Li, J., Zhang, W., Xiao, C., Zhang, S., Nian, C., Li, J., Su, D., Chen, L., Zhao, Q., et al. (2021). Glycogen accumulation and phase separation drives liver tumor initiation. Cell 184, 5559–5576.e19.

19. Tang, K., Zhu, L., Chen, J., Wang, D., Zeng, L., Chen, C., Tang, L., Zhou, L., Wei, K., Zhou, Y., et al. (2021). Hypoxia promotes breast cancer cell growth by activating a glycogen metabolic program. Cancer Res. 81, 4949–4963.

20. Iida, Y., Aoki, K., Asakura, T., Ueda, K., Yanaihara, N., Takakura, S., Yamada, K., Okamoto, A., Tanaka, T., and Ohkawa, K. (2012). Hypoxia promotes glycogen synthesis and accumulation in human ovarian clear cell carcinoma. Int. J. Oncol. 40, 2122–2130.

21. Young, L.E.A., Conroy, L.R., Clarke, H.A., Hawkinson, T.R., Bolton, K.E., Sanders, W.C., Chang, J.E., Webb, M.B., Alilain, W.J., Vander Kooi, C.W., et al. (2022). In situ mass spectrometry imaging reveals heterogeneous glycogen stores in human normal and cancerous tissues. EMBO Mol. Med. 14, e16029.

22. Favaro, E., Bensaad, K., Chong, M.G., Tennant, D.A., Ferguson, D.J., Snell, C., Steers, G., Turley, H., Li, J.L., Günther, U.L., et al. (2012). Glucose utilization via glycogen phosphorylase sustains proliferation and prevents premature senescence in cancer cells. Cell Metab. 16, 751–764.

23. Ji, Q., Li, H., Cai, Z., Yuan, X., Pu, X., Huang, Y., Fu, S., Chu, L., Jiang, C., Xue, J., et al. (2023). PYGL-mediated glucose metabolism reprogramming promotes EMT phenotype and metastasis of pancreatic cancer. Int. J. Biol. Sci. 19, 1894–1909.

24. Heden, T.D., Chow, L.S., Hughey, C.C., and Mashek, D.G. (2022). Regulation and role of glycophagy in skeletal muscle energy metabolism. Autophagy 18, 1078–1089.

25. Koutsifeli, P., Varma, U., Daniels, L.J., Annandale, M., Li, X., Neale, J.P.H., Hayes, S., Weeks, K.L., James, S., Delbridge, L.M.D., et al. (2022). Glycogen-autophagy: Molecular machinery and cellular mechanisms of glycophagy. J. Biol. Chem. 298, 102093.

26. Han, Z., Zhang, W., Ning, W., Wang, C., Deng, W., Li, Z., Shang, Z., Shen, X., Liu, X., Baba, O., et al. (2021). Model-based analysis uncovers mutations altering autophagy selectivity in human cancer. Nat. Commun. 12, 3258.

27. Reeves, S.R., and Gozal, D. (2004). Platelet-activating factor receptor modulates respiratory adaptation to long-term intermittent hypoxia in mice. Am. J. Physiol. Regul. Integr. Comp. Physiol. 287, R369–R374.

28. Drummond, S.E., Burns, D.P., O’Connor, K.M., Clarke, G., and O’Halloran, K.D. (2021). The role of NADPH oxidase in chronic intermittent hypoxia-induced respiratory plasticity in adult male mice. Respir. Physiol. Neurobiol. 292, 103713.

29. Jackson, E.L., Willis, N., Mercer, K., Bronson, R.T., Crowley, D., Montoya, R., Jacks, T., and Tuveson, D.A. (2001). Analysis of lung tumor initiation and progression using conditional expression of oncogenic K-ras. Genes Dev. 15, 3243–3248.

30. Jackson, E.L., Olive, K.P., Tuveson, D.A., Bronson, R., Crowley, D., Brown, M., and Jacks, T. (2005). The differential effects of mutant p53 alleles on advanced murine lung cancer. Cancer Res. 65, 10280–10288.

31. Conroy, L.R., Chang, J.E., Sun, Q., Clarke, H.A., Buoncristiani, M.D., Young, L.E.A., McDonald, R.J., Liu, J., Gentry, M.S., Allison, D.B., et al. (2022). High-dimensionality reduction clustering of complex carbohydrates to study lung cancer metabolic heterogeneity. Adv. Cancer Res. 154, 227–251.

32. van der Ploeg, A.T., and Reuser, A.J.J. (2008). Pompe’s disease. Lancet 372, 1342–1353.

33. Conroy, L.R., Clarke, H.A., Allison, D.B., Valenca, S.S., Sun, Q., Hawkinson, T.R., Young, L.E.A., Ferreira, J.E., Hammonds, A.V., Dunne, J.B., et al. (2023). Spatial metabolomics reveals glycogen as an actionable target for pulmonary fibrosis. Nat. Commun. 14, 2759.

34. Johnson, L.N. (1992). Glycogen phosphorylase: control by phosphorylation and allosteric effectors. FASEB J. 6, 2274–2282.

35. Ren, J.M., and Hultman, E. (1990). Regulation of phosphorylase a activity in human skeletal muscle. J. Appl. Physiol. 69, 919–923.

36. Sun, R.C., Dukhande, V.V., Zhou, Z., Young, L.E.A., Emanuelle, S., Brainson, C.F., and Gentry, M.S. (2019). Nuclear glycogenolysis modulates histone acetylation in human non-small cell lung cancers. Cell Metab. 30, 903–916.

37. Hakim, F., Wang, Y., Zhang, S.X., Zheng, J., Yolcu, E.S., Carreras, A., Khalyfa, A., Shirwan, H., Almendros, I., and Gozal, D. (2014). Fragmented sleep accelerates tumor growth and progression through recruitment of tumor-associated macrophages and TLR4 signaling. Cancer Res. 74, 1329–1337.

38. Nanduri, J., Yuan, G., Kumar, G.K., Semenza, G.L., and Prabhakar, N.R. (2008). Transcriptional responses to intermittent hypoxia. Respir. Physiol. Neurobiol. 164, 277–281.

39. Cubillos-Zapata, C., Balbás-García, C., Avendaño-Ortiz, J., Toledano, V., Torres, M., Almendros, I., Casitas, R., Zamarrón, E., García-Sánchez, A., Feliu, J., et al. (2019). Age-dependent hypoxia-induced PD-L1 upregulation in patients with obstructive sleep apnoea. Respirology 24, 684–692.

40. Liu, Y., Lu, M., Chen, J., Li, S., Deng, Y., Yang, S., Ou, Q., Li, J., Gao, P., Luo, Z., et al. (2022). Extracellular vesicles derived from lung cancer cells exposed to intermittent hypoxia upregulate programmed death ligand 1 expression in macrophages. Sleep Breath. 26, 893–906.

